# Single-cell RNA sequencing reveals cell-type specific *cis*-eQTLs in peripheral blood mononuclear cells

**DOI:** 10.1101/177568

**Authors:** M.G.P. van der Wijst, H. Brugge, D.H. de Vries, L.H. Franke

**Affiliations:** Department of Genetics, University of Groningen, University Medical Center Groningen, Groningen, The Netherlands

## Abstract

Most disease-associated genetic risk factors are regulatory. Here, we generated single-cell RNA-seq data of ∼25,000 peripheral blood mononuclear cells from 45 donors to identify how genetic variants affect gene expression. We validated this approach by replicating previously published whole blood RNA-seq *cis-*expression quantitative trait loci effects (*cis-*eQTLs), but also identified new cell type-specific *cis-*eQTLs. These eQTLs give additional insight into the downstream consequences of genetic risk factors for immune-mediated diseases.

## Main text

Genome-wide association studies identified thousands of genetic variants (single nucleotide polymorphisms, SNPs) that increase disease risk or are associated with a specific trait.^1^ Most of these SNPs have only a small effect on disease risk. Moreover, this type of analysis holds no information on which genes, and subsequently, which pathways are involved. In contrast, we and others have previously shown that these SNPs can have a large effect on gene expression in blood, and by correlating these so-called expression quantitative trait loci (eQTLs), insights can be gained in the downstream consequences of disease.^2^

Based on previous studies^3-5^, we hypothesize that the downstream expression effects of many disease risk-SNPs are cell type-specific. Previously, purified cell types^4, 6-8^ or deconvolution methods^9, 10^ have been used to identify cell type-specific eQTLs. However, these methods are either biased (cell-sorting), or are of limited use for less abundant cell types and dependent on accurate cell type quantification by previously defined marker genes (deconvolution). In contrast, single-cell RNA sequencing (scRNA-seq) could enable identification of cell type-specific eQTLs using an unbiased approach and can be used to investigate rare cell types^11^.

To determine the cell type-specific effects of genetic variation on gene expression, an eQTL analysis was performed in ∼25,000 peripheral blood mononuclear cells (PBMCs) from 45 donors of the population cohort Lifelines Deep^12^. However, before executing a genome-wide eQTL analysis, we first validated our approach by focusing only on previously reported top *cis*-eQTLs detected in whole blood DeepSAGE^13^ (which is also a 3’-end oriented RNA-sequencing strategy) or bulk RNA-seq^14^ data. For this analysis, we treated our data as being bulk PBMC data, from now on referred to as “total PBMCs”. We could significantly replicate 50 and 311 *cis*-eQTLs (of which 96.0% and 90.4% had the same allelic direction) for the DeepSAGE and RNA-seq study, respectively (Fig. 1a, Suppl. Table 1). The few disconcordant eQTLs between the datasets may be partly related to the slightly different composition of whole blood versus PBMCs, i.e. whole blood contains some extra cell types, including neutrophils and eosinophils. Altogether, this indicated that *cis*-eQTL mapping using scRNA-seq data yields reliable results.

**Table 1.**
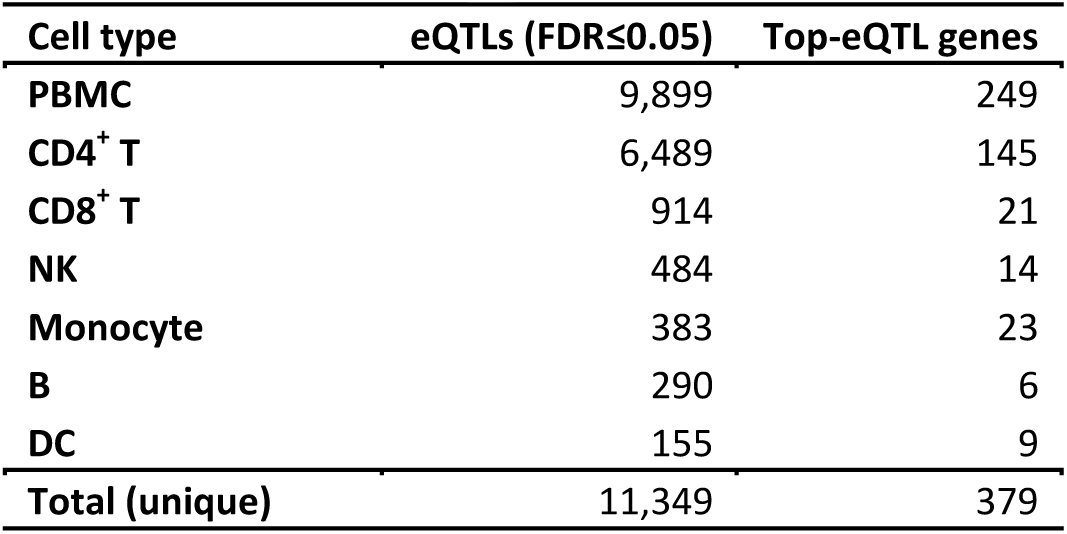
**eQTLs identified per cell type**

**Figure 1.**
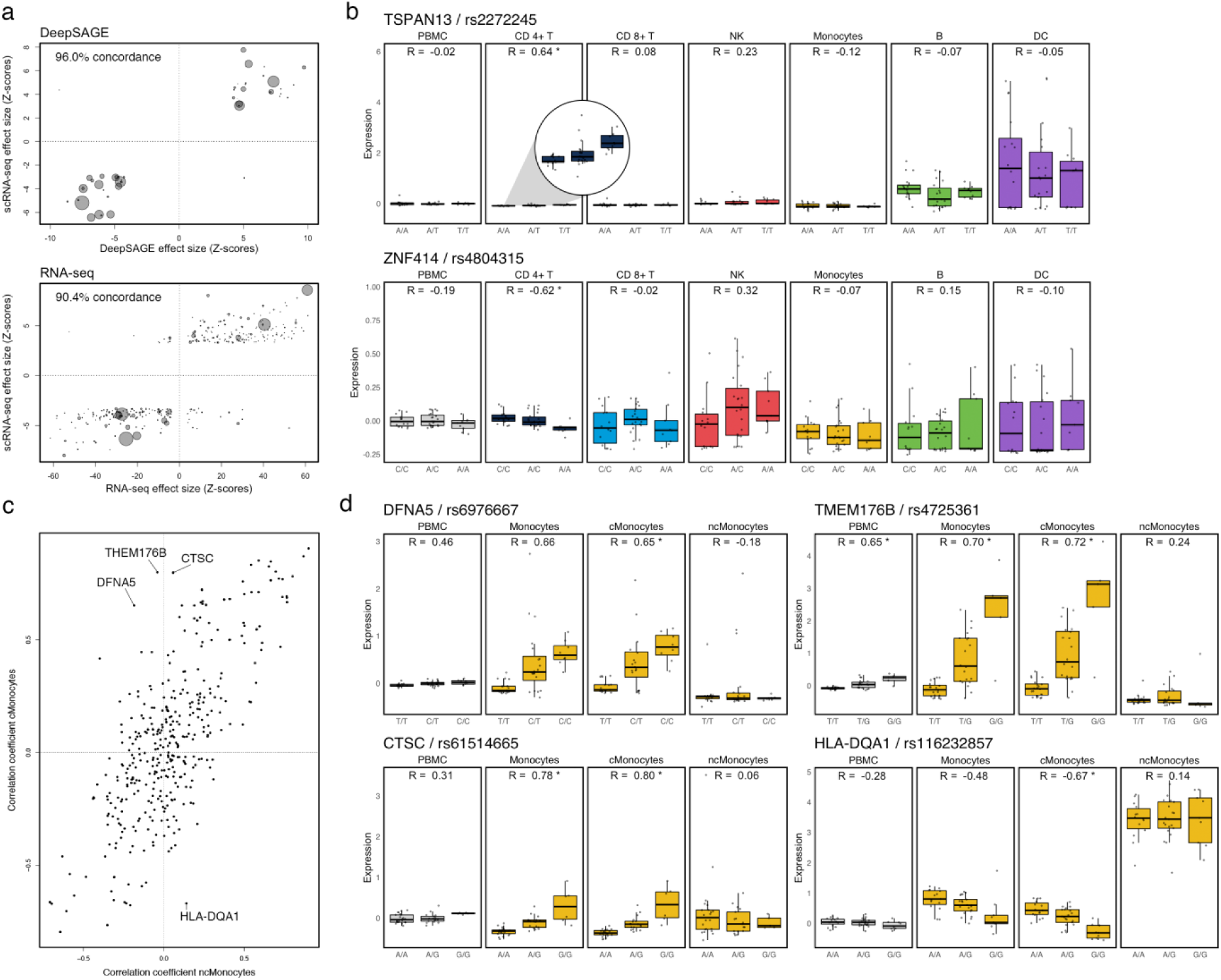
*cis*-eQTL analysis in single-cell RNA-seq data. (**a**) Concordance between total PBMC population scRNA-seq eQTLs and (top) whole blood DeepSAGE (3’-end transcript reads) or (bottom) bulk RNA-seq data. The size of each dot represents the mean expression of the *cis*-regulated gene in the whole dataset. (**b**) Examples of undetectable eQTLs in the total PBMC population caused by (top) masking of the eQTL present in CD4^+^ T cells but absent in DCs with comparatively high expression of the *cis*-regulated gene or (bottom) opposite allelic effects in CD4^+^ T and NK cells. (**c**) Correlation coefficient of each top SNP-gene combination for the cMonocytes against the ncMonocytes. (**d**) The 4 outliers highlighted in **c** visualized: eQTLs specifically affecting expression in the cMonocytes, and not the ncMonocytes. Each dot represents a donor. Box plots show the median, the first and third quartiles, and 1.5 times the interquartile range. R, correlation coefficient; *FDR≤0.05.

We then performed a genome-wide *cis-*eQTL analysis on the total PBMCs and each of the major cell types we had identified (Suppl. Fig. 1a, 1b). This was done by averaging the normalized gene expression of all cells per cell type per donor for the following cell types: CD4^+^ T cells, CD8^+^ T cells, natural killer (NK) cells, monocytes, B cells and dendritic cells (DC). In total, 379 unique top-eQTLs were detected (false discovery rate (FDR)≤0.05), of which 249 were found to be top-eQTLs in the total PBMCs (Table 1). Of these 249, we could replicate 181 (90.1% concordance) in whole blood RNA-seq data^14^ (Suppl. Table 2). Within the 130 remaining top-eQTLs, only 48 were not found significant in total PBMCs, but only in certain cell types (Suppl. Table 2).

Several factors can explain why not all *cis*-eQTLs can be reproduced in the total PBMCs. First, unless using a very large sample size, the signal of an eQTL in a less abundant cell type may be diluted out in the total PBMCs. Along these lines, when using 2,116 whole blood RNA-seq samples from Zhernakova et al., we could replicate 29 out of 48 eQTLs (100% concordance). Leaving us with 19 *cis*-eQTLs that could not be found using such a large sample size of whole blood RNA-seq data. Secondly, in a common cell type, the eQTL may be masked by very high expression of this *cis*-regulated gene in a less abundant cell type lacking this eQTL. We observed this situation in CD4^+^ T-cells, where rs2272245 significantly affected the expression of the very lowly expressed *TSPAN13* gene in *cis*. However, this effect was neither observed in any of the other cell types, nor in the total PBMCs, as in the total PBMCs the very high *TSPAN13* expression of DCs masked the eQTL found in CD4^+^ T-cells (Fig. 1b). Finally, eQTLs may show opposite allelic effects across different cell types. The small sample size of the less abundant cell types (Suppl. Fig. 1c) did not allow for the identification of two significant opposite allelic effects. Despite this, in CD4^+^ T cells, the A allele of rs4804315 significantly decreased the expression of *ZNF414* in *cis* (Nominal P-value = 6.09*10^−6^), whereas in NK cells this same allele increased expression of *ZNF414* (Nominal P-value = 0.0339) (Fig. 1b). However, it cannot be excluded that specifically in NK cells, the effect of rs4804315 on *ZNF414* expression is the result of a residual effect on *ZNF414* expression of a second independent eQTL.

All the above-mentioned factors that may influence the reproducibility of *cis*-eQTLs in total PBMCs can be overcome using RNA-seq data of purified cell types. Indeed, 4 out of the 19 remaining *cis*-eQTLs (100% concordance) were reproduced in purified RNA-seq data of the Blueprint consortium (naïve CD4^+^ T cells and CD14^+^ monocytes)^15^ or Kasela et al. (CD4^+^ and CD8^+^ T cells)^6^ (Suppl. Table 3). Hence, only a small fraction (15 cis-eQTLs) of eQTLs were not identified before using RNA-seq data of either whole blood or purified cells (CD4^+^ and CD8^+^ T cells or monocytes). Even though some *cis*-eQTLs were only found significant in specific cell types, this does not imply they are cell type-specific; less abundant cell types may lack the statistical power required to detect these *cis*-eQTLs. Instead of tissue-by-tissue eQTL analysis, meta-analysis may increase the power to detect effects particularly in the less abundant cell types. However, commonly used meta-analyses for bulk RNA-seq, such as eQTL-BMA^16^ or Meta-Tissue^17^, are computationally too demanding for large scRNA-seq data or do not define the cell type in which the eQTL effect is present.^17, 18^

The main advantage of scRNA-seq data for eQTL analysis is the flexibility by which any cell population of interest can be selected for analysis. In contrast, when using RNA-seq data of purified cell types, one cannot retrieve data from subcell types anymore. Moreover, the finer differences between subcell types may not always be recapitulated by different cell membrane markers, whereas they are on the gene expression level. To show the added value of performing an eQTL analysis on scRNA-seq data, we performed an analysis on the two identified subtypes of monocytes: classical (cMonocytes) and non-classical monocytes (ncMonocytes). When plotting the correlation coefficient of each top SNP-gene combination for the cMonocytes against the ncMonocytes, we revealed 4 clear examples in which we could pinpoint the eQTL effect specifically to the cMonocyte subset (Fig. 1c). Each of these 4 eQTLs were already previously identified in RNA-seq data of purified CD14^+^ monocytes^15^, but by using the scRNA-seq data we could now identify these effects to be specific for the cMonocyte subset (Fig. 1d).

The above examples clearly show the benefit of scRNA-seq data for eQTL analysis, but we expect scRNA-seq data to offer many other opportunities for selecting cells of interest for eQTL analysis. For example, one could use the intercellular variation within scRNA-seq data to group cells along the cell cycle, along a differentiation path or along a response to an environmental stimulus. By doing so, one may be able to define eQTLs that are amplified or abrogated depending on the cell cycle phase, differentiation status or environmental stimulus.

In the future, it is essential to improve the statistical power to detect eQTLs in scRNA-seq data. Especially the less abundant, less well-studied cell types are expected to benefit from this. One could do so by overcoming the zero-inflated expression that is inherent to scRNA-seq data by clever computational strategies that recover some of the expression^19^ or by detecting eQTL effects on gene expression networks instead of individual genes^20^, thereby improving the reliability of the gene expression measurements. In addition, combining multiple scRNA-seq studies in one big meta-analysis is expected to exponentially increase the number of eQTLs that will be found.

In conclusion, this proof-of-concept study shows the feasibility of scRNA-seq data for eQTL analysis. The identified top-eQTLs replicated well with earlier reported whole blood RNA-seq data. Moreover, we extended the list of genes that are known to be under genetic regulation or specified the cell type in which the effect is most prominent. We expect that this study is just the start of many new opportunities to study how genetic variation is affecting single-cell gene expression levels.

## Acknowledgements

We are very grateful to all the volunteers who participated in this study. Moreover, we thank Jackie Dekens for arranging informed consent and contact with LifeLines. This study was supported by an ERC starting grant (grant #637640).

## Author contribution

MW generated the scRNA-seq data. MW, HB and DV performed bioinformatics and statistical analyses. MW and LF designed the study and wrote the manuscript. All authors discussed the results and commented on the manuscript.

## Ethics approval and consent to participate

The LifeLines DEEP study was approved by the ethics committee of the University Medical Centre Groningen, document number METC UMCG LLDEEP: M12.113965. All participants signed an informed consent form prior to study enrollment. All procedures performed in studies involving human participants were in accordance with the ethical standards of the institutional and/or national research committee and with the 1964 Helsinki declaration and its later amendments or comparable ethical standards.

